# Developmental pathways regulate cytokine-driven effector and feedback responses in the intestinal epithelium

**DOI:** 10.1101/2020.06.19.160747

**Authors:** Håvard T. Lindholm, Naveen Parmar, Claire Drurey, Jenny Ostrop, Alberto Díez-Sanchez, Rick M. Maizels, Menno J. Oudhoff

## Abstract

The intestinal tract is a common site for infection, and it relies on an appropriate immune response to defend against pathogens. The intestinal epithelium has an important role in effector responses, which is coordinated by immune-type specific cytokines. It is incompletely understood how cytokines drive epithelial responses. Here, using organoid-cytokine co-cultures, we provide a comprehensive analysis of how key cytokines affect the intestinal epithelium, and relate this to *in vivo* infection models. We use imaging, based on a convolutional neural network, and transcriptomic analysis to reveal that cytokines use developmental pathways to define intestinal epithelial effector responses. For example, we find IL-22 and IL-13 dichotomously induce goblet cells, in which only IL-13 driven goblet cells are associated with NOTCH signaling. We further show that IL-13 induces BMP signalling to act as a negative feedback loop in IL-13 induced tuft cell hyperplasia, an important aspect of type 2 immunity. Together, we show that targeting developmental pathways may be a useful tool to tailor epithelial effector responses that are necessary for immunity to infection.

## Introduction

Infections of the gastrointestinal tract remain one of the leading health issues worldwide (Kirk et al. 2015). The ability to clear infections relies on the capacity of the immune system to mount an appropriate response. Immune responses can be divided into 3 types. Broadly; type 1 responses are launched against intracellular pathogens such as bacteria and viruses and are associated with the cytokine interferon gamma (IFNγ), type 2 responses are generated to fight off parasitic infections and are driven by interleukin (IL)-4 and IL-13, and finally, type 3 responses are mounted against extracellular pathogens such as bacteria and are associated with IL-22 and IL-17 (Kara et al. 2014; Raphael et al. 2015).

In addition to defining the immune landscape, cytokines directly affect intestinal epithelial cells (IECs) to drive immune-type specific effector responses (Biton et al. 2018). The intestinal epithelium consists of a single layer of cells and is responsible both for taking up nutrients as well as providing a protective barrier. One of the hallmarks of IECs is their rapid turnover (3-5 days), which allows for prompt cellular responses, for example, to expand goblet cells which can produce protective mucus. This plasticity of IECs makes them particularly well-suited to defend against pathogens (Tetteh, Farin, and Clevers 2015). In addition to responding to immune cues, the epithelium can also be involved in tailoring immune responses. For example, tuft cells, which are important for defending against parasitic helminths, are the main source of IL-25 to control ILC2 populations both in homeostasis and upon helminth infection (Gerbe et al. 2016; Howitt et al. 2016; Moltke et al. 2016). As tuft cells rapidly expand upon exposure to type 2 cytokines, they exemplify both how the intestinal epithelium changes upon an immune response and how it can partake in shaping it.

Intestinal organoids are self-organizing structures that are useful to study epithelial (stem) cell biology (Dedhia et al. 2016). They are particularly instructive to identify epithelial-intrinsic responses as they lack any other cell type normally present *in vivo*, such as fibroblasts. However, organoid cultures also come with challenges as most metrics cannot capture the complexity of these large multicellular structures. To capture this complexity, a recent study used single cell RNA sequencing (scRNA-seq) to systemically compare single cell responses to cytokines of the three immune environments (Biton et al. 2018). However, there are still many unclear aspects to how cytokines induce epithelial responses, especially mechanistically. Here we combine quantitative imaging with bulk RNA-seq to define how cytokines intercept developmental pathways to instruct epithelial differentiation and maturation. Most prominently, we identify a feedback loop by which IL-13 induced tuft cell hyperplasia is self-limiting in a BMP-dependent manner.

## Results

### IFNγ, IL-13, and IL-22 uniquely affect organoid growth and morphology as determined using a convolutional neural network-based classification

Intestinal organoids grow as a morphologically heterogeneous population, where a fraction of organoids grow as immature “spheroids” and the rest form mature “budding” organoids. The change in distribution of these morphological categories has been used to study diverse processes in intestinal organoid biology (Mustata et al. 2013; Schwank et al. 2013), but currently there are limited options available for systematically comparing these morpho-logical groups. Building on our recent work on organoid segmentation (Ostrop et al. 2020), we here developed an automated image analysis pipeline that segments organoid objects from the background and subsequently classifies them into “spheroid” or “budding” categories based on a convolutional neural network (Fig. 1A). An optional manual correction step allowed for inclusion of organoids that were difficult to automatically segment or classify otherwise (Fig. S1A). Comparing manually verified with automatically classified and segmented images had a good correlation in analysis of >20,000 organoids (Fig. S1B). Furthermore, we find that in time the number of spheroids decrease and appear darker in appearance, confirming that our systematic approach captures what is observed visually (Fig. 1B,C and S1D).

**Figure 1:**
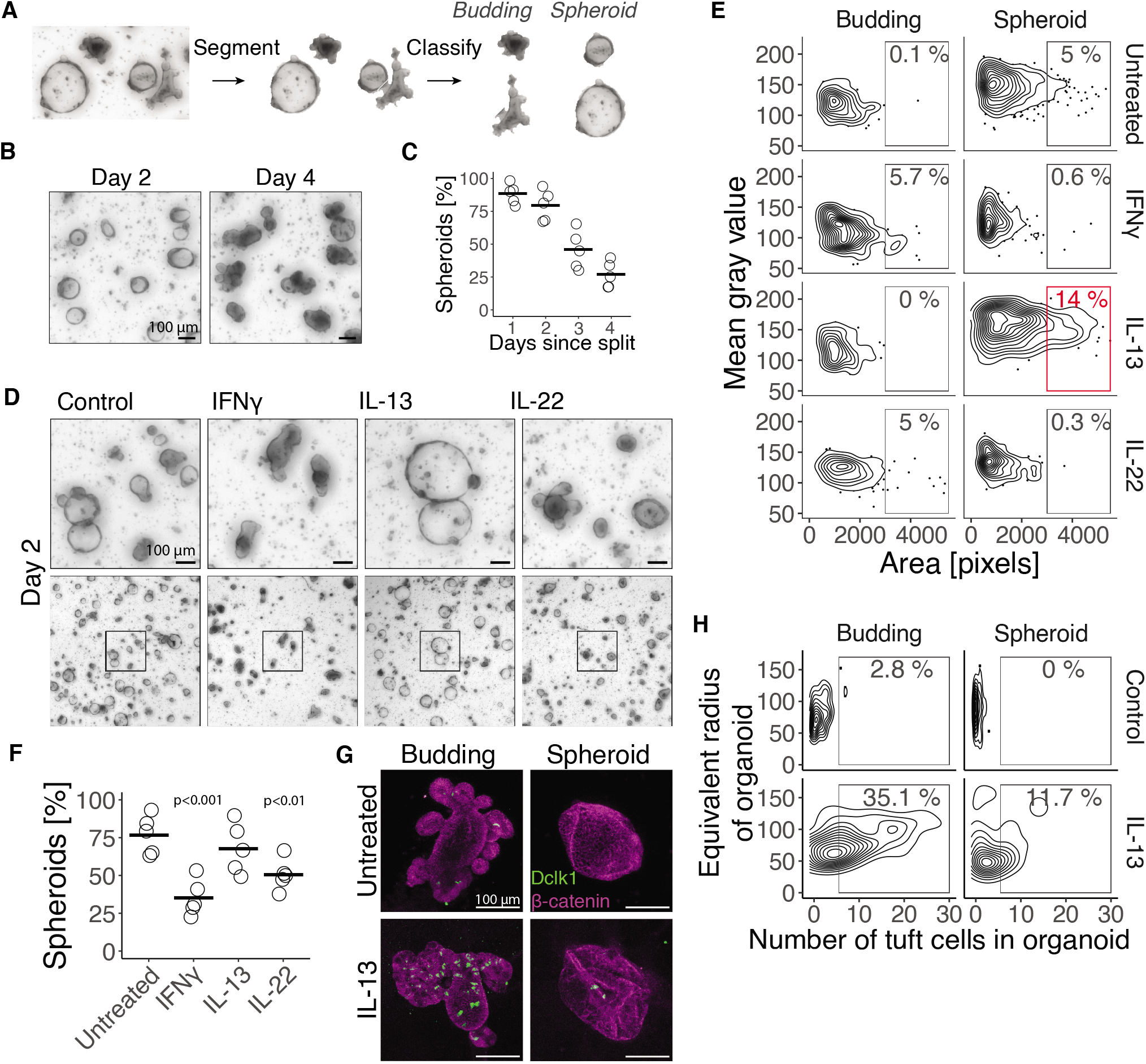
Cytokines change organoid growth and morphology. **A**, Image segmentation and classification of organoid images. **B**, Representative images of untreated organoids. **C**, Average fraction of spheroids in each image in same experiment as B. Each circle is one biological replicate and is the average of at least 3 images and in total this plot represents 16,462 organoids. **D**, Bright field images of small intestinal organoids treated with 10 ng/mL cytokine since the day of splitting. **E**, Distribution of gray value (higher value is whiter) and area of all organoids in same experiment as D. Percentages are relative to total number of organoids in that treatment. Plot shows distribution of in total 2,310 organoids, representative result of 5 mice. **F**, Average fraction of spheroids at day 2 since splitting. Each circle is one biological replicate and is the average of at least 3 images and in total represents 10,462 organoids. Statistics calculated with a two-tailed T-test on mean of biological replicates compared to untreated. **G**, Representative confocal images of small intestinal organoids at day 3 since splitting. **H**, Distribution of number of tuft cells and size of all organoids from same experiment as G. Percentages are relative to total number of organoids in that treatment. Plot shows the distribution of 192 organoids, representative in 4 experiments of 4 mice. All images are projections of Z-stacks.

To mimic specific immune responses, we treated organoids with IFNγ, IL-13, or IL-22 to represent type 1, 2, and 3 responses respectively. As was previously found by other groups (Farin et al. 2014; Lindemans et al. 2015), long term IL-22 or IFNγ treatment ultimately leads to organoid disintegration (Fig. S1C). At an earlier time point (day 2), this is characterized by a darker appearance, reduced percentage of spheroids, and an increased percentage of budding organoids (Fig. 1D-F). In contrast, a population of IL-13 treated organoids form large spheroids at day 2 (Fig. 1D and 1E). Of note, the differentiation into mature epithelial lineages commonly occurs in budding organoids (Serra et al. 2019).

Tuft cells are a specific epithelial cell type involved in type 2 immunity, and induced by IL-13 (Gerbe et al. 2016; Moltke et al. 2016). We next combined our classification setup with confocal imaging to automatically quantify the number of tuft cells in organoid subtypes. This configuration allows for classification and counting of tuft cells in hundreds of organoids and supports both automatic estimates of tuft cell number and a more accurate manually curated count (Fig. S1E). Indeed, we find that tuft cells primarily appear in budding organoids both in control and IL-13-treated conditions (Fig. 1G,H). Together, these data show that classification of organoids provides a tractable measure of epithelial responses such as immune cues.

### IFNγ, IL-13, and IL-22 uniquely affect RNA expression in organoids in a manner aligned to *in vivo* infection profiles

To assess in detail how IFNγ, IL-13, and IL-22 affect intestinal epithelial cells we performed bulk RNA-seq on organoids treated with indicated cytokines for 24 hours compared to untreated controls. In a PCA plot, all four conditions grouped separately, highlighting the different effector responses required for each type of immunity (Fig. 2A). This is further exemplified by uniquely induced genes for each condition (Fig. 2B). Furthermore, IL-22 and IL-13 induced genes associated with various cellular responses as determined by GO term analysis (Fig. S2A,B). In contrast, IFNγ induced genes associated primarily with antigen presentation, as the most significant GO terms were related to this (Fig. 2C). In support, flow cytometry analysis of IFNγ treated organoids showed that all cells gain MHC-II and 90% gained MHC-I expression (Fig. 2D). *In vivo*, during homeostasis, specifically Lgr5+ cells express MHC-II (Biton et al. 2018). Our data here thus suggests that during infection all epithelial cell types may be involved in altering the local immune cell environment through antigen presentation.

**Figure 2:**
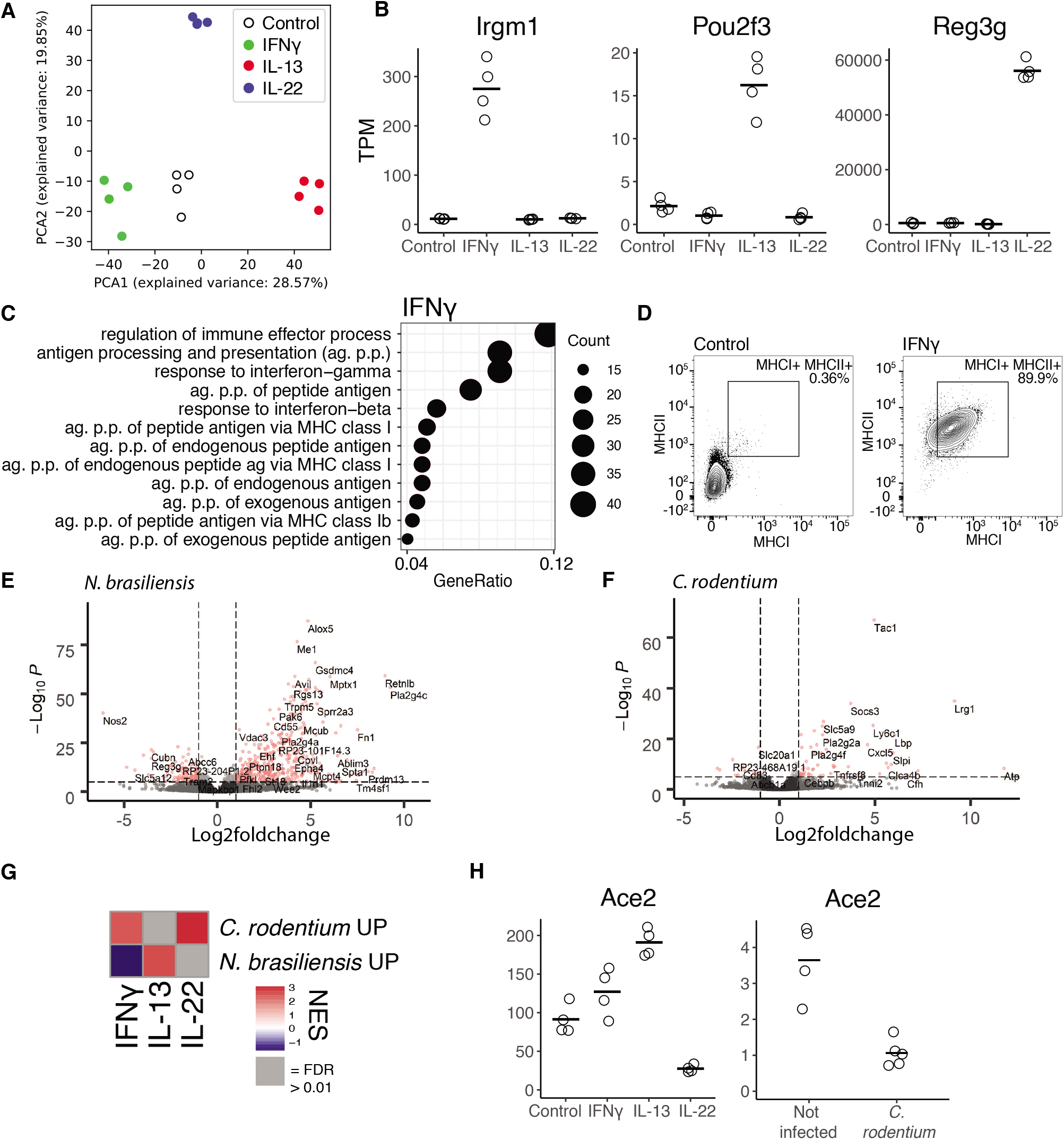
Cytokines modify development of small intestinal epithelium in a cytokine-specific manner and correspond to *in vivo* infection models. **A**, PCA plot of log2(TPM + 1) values determined with RNAseq of small intestinal organoids treated with 10 ng/mL cytokines for 24 hours. Four biological replicates per treatment. **B**, Expression of genes from same data as A. Adjusted p-value relative to control: *Irgm1*: IFNγ: < 10^−163^, IL-13: NS, IL-22: < 0.01, *Pou2f3*: IFNγ: < 0.05, IL-13: < 10^−35^, IL-22: < 0.05, *Reg3g*: IFNγ: NS, IL-13: < 0.05 and IL-22: < 10^−75^. **C**, Top 12 enriched biological process GO-terms of genes significantly upregulated in small intestinal organoids treated with IFNγ compared to control, p-adjusted < 0.01 and log2fc > 1. **D**, Flow cytometry of small intestinal organoids treated for 4 days with cytokine, representative plot of 4 mice. **D,E**, Volcano plot of RNAseq from intestinal epithelial tissue extracted from mice infected with indicated pathogen. **G**, GSEA of the top 300 most significantly up-regulated genes from intestinal epithelium from mice infected with indicated organism, see supplementary file 1. This is compared to intestinal organoids stimulated with indicated cytokine compared to control. **H**, See B. Adjusted p-value relative to relevant control: IFNγ: < 0.15, IL-13: > 10^−10^, IL-22: > 10^−13^ and *C. rodentium*: > 10^−7^. NES = normalised enrichment score, TPM = transcripts per million and NS = not significant.

To assess the biological relevance of our *in vitro* data, we performed RNA-seq on intestinal epithelium from mice infected with *Nippostrongylus brasiliensis* or *Citrobacter rodentium*, which are classical infection models inducing type 2 and 3 immune responses respectively (Fig. 2E,F, S2C,D). We found that epithelium isolated from *C. rodentium* infected mice aligned with IL-22- and IFNγ-treated organoids using gene set enrichment analysis (GSEA) (Fig. 2G). In contrast, epithelium isolated from *N. brasiliensis* infected mice was similar to IL-13 treated-organoids (Fig. 2G). This thus aligns with the standard model of type 2 and type 3 immune responses and indicates that organoid-cytokine co-cultures provide a relevant model to replicate *in vivo* epithelial responses.

Epithelium is not only is important in immune effector responses, but can also act as a port of entry for pathogens, for example, SARS-CoV-2 relies on epithelial-expressed ACE2. Thus, given the current COVID-19 pandemic, we were interested whether cytokines could affect *Ace2.* This could be relevant in relation to previous infections or co-infection. In line with other work, we find a slight increase of *Ace2* in IFN-g treated organoid (Ziegler et al. 2020) (Fig. 2H). In addition, we find a stronger increase of *Ace2* upon IL-13 treatment. In stark contrast, we find IL-22 greatly reduces *Ace2* expression *in vitro* and *in vivo* after *C. rodentium* infection (Fig. 2H). As Citro is associated with both IL-22 and IFNγ, we conclude that the IL-22 mediated repression is dominant over IFNγ related activation.

### Cytokines uniquely affect the intestinal epithelial cellular composition

To test the effect of IFNγ, IL-13 and IL-22 on cell lineage differentiation, we used cell-type specific gene signatures acquired through scRNAseq (Haber et al. 2017). GSEA revealed that these cytokines broadly affect intestinal cell lineage differentiation in an expected manner (Fig. 3A). For example, giving IL-13 to organoids induced a tuft cell signature, supported by tuft cell staining *in vitro* (Fig. 1G) and *in vivo* upon *N. brasiliensis* infection (Fig. S3A-C). In addition, both IL-13 and IL-22 induced a goblet-cell associated gene signature (Fig. 3A). However, close examination showed that each cytokine induced a different set of goblet-cell genes with relatively little overlap both after 24 and 72 hours of cytokine stimulation (Fig. 3B, S4A). This is exemplified by the traditional goblet cell marker *Muc2* being only induced by IL-13 whereas RELM-beta *(Retnlb)* was induced more strongly by IL-22 (Fig. 3C,D, S4B). Confocal staining confirms that MUC2 was only induced by IL-13, and that RELM-beta was more strongly induced by IL-22 where it leads to accumulation of smaller cells that we presume to be MUC2 negative (Fig. 3E,F). Canonical induction of goblet cells occurs through inhibition of NOTCH and further relies on ATOH1 and SPDEF (Noah et al, Shroyer, Exp Cell Res 2009). Indeed, IL-13 induced *Atoh1* and *Spdef*, however, IL-22 did not, indicating that IL-22 induces goblet cells in a non-canonical manner (3G,H, S4,C). GO-term analysis of the goblet cell genes uniquely induced by IL-13 and IL-22 revealed different terms, for example, “response to endoplasmatic stress” was the top GO term associated with goblet cell genes uniquely induced by IL-22 (Fig. 3I, S4D,E). Together, this indicates that IL-13 and IL-22 likely induce different types of goblet cells.

**Figure 3:**
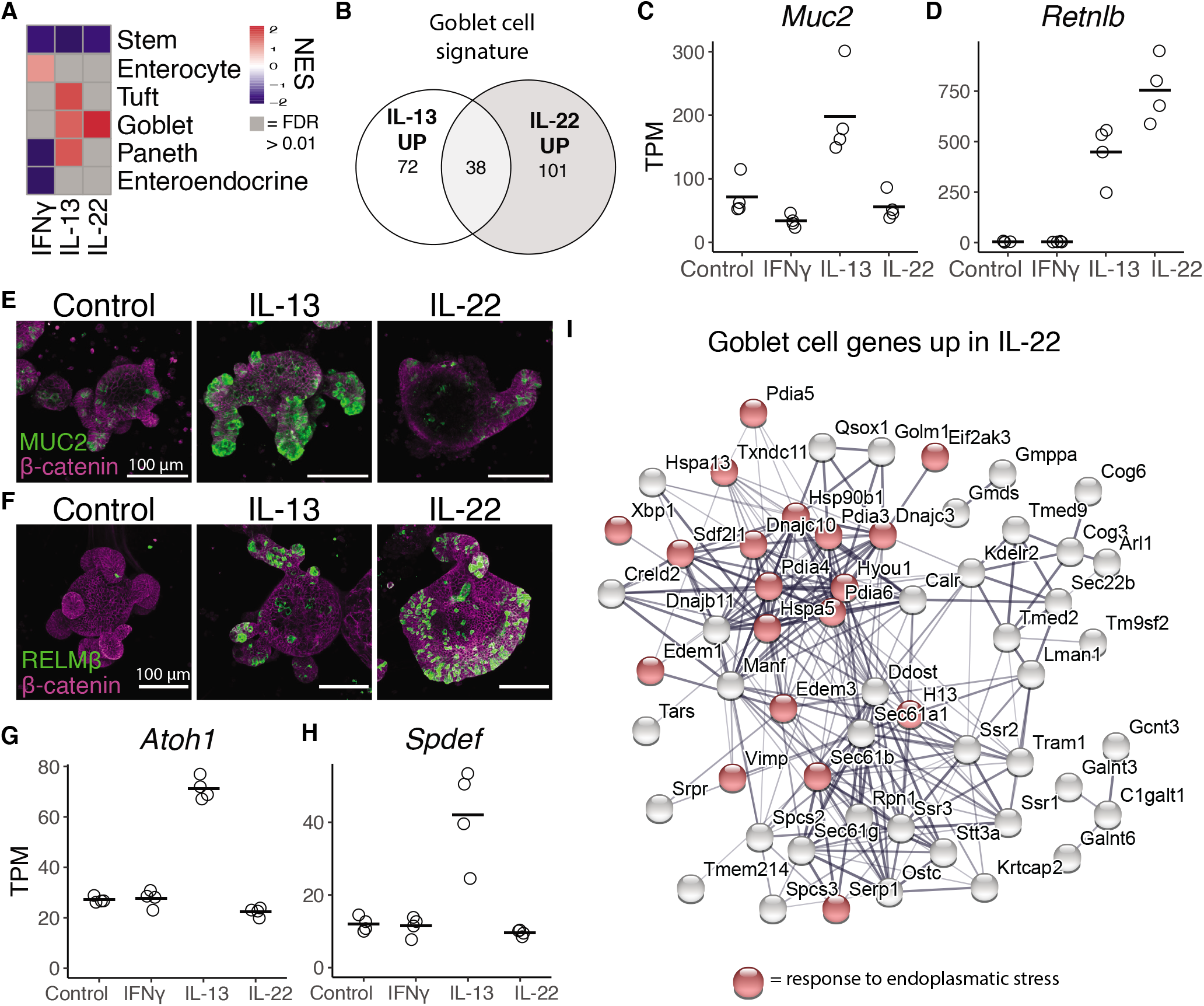
IL-13 and IL-22 induce different subsets of goblet cell genes. **A**, Heatmap of NES values from GSEA analysis of gene sets representing different cell types, see supplementary file 1 for gene sets. **B**, Distribution of how the goblet cell gene set from plate based scRNAseq from Haber et al is changed upon 24 hour IL-13 and IL-22 treatment. Up is defined as lg2fc>0.5 and p-adj<0.05. **C,D**, Gene expression from intestinal organoids treated for 24 hours with indicated cytokine. Black bar is mean. Adjusted p-values relative to control: *Muc2*: IFNγ: <10^−4^, IL-13: <10^−8^ and IL-22: NS. *Retnlb*: IFNγ: NS, IL-13: <10^−43^ and IL-22: = <10^−63^. **E,F**, Confocal staining of MUC2 at day 2 (E) and RELMB at day 4 (F) in small intestinal organoids. Representative images of three (E) and two (F) mice in independent experiments. **G, H**, See C. Adjusted p-values relative to control: *Atho1*: IFNγ: NS, IL-13: <10^−32^ and IL-22: NS. *Spdef*: IFNγ: NS, IL-13: <10^−15^ and IL-22: NS. **I**, Protein network from the STRING database of goblet cell genes that are up-regulated upon IL-22 treatment, but not upon IL-13 treatment. Red nodes are found in the biological process GO-term “response to endoplasmatic reticulum”. NES = normalised enrichment score, TPM = transcripts per million and NS = not significant. All images are projections of Z-stacks.

### Cytokines utilize developmental signaling pathways to regulate effector responses

We were interested in how cytokine treatment leads to cellular effector responses. Intestinal epithelium relies on NOTCH-, WNT-, BMP- and HIPPO-signaling to maintain homeostatic differentiation of cell lineages. We hypothesized that cytokines may use these developmental pathways in directing epithelial cell differentiation. We generated gene sets for these different pathways from published transcriptome datasets (Gregorieff et al. 2015; Kim et al. 2014; Qi et al. 2017; Yin et al. 2014). GSEA analysis revealed that cytokines, and in particular IL-13 and IL-22, alter transcription of genes normally associated with HIPPO, NOTCH, and BMP pathways (Fig. 4A). Specifically, the HIPPO signature was enriched in both IL-13 and IL-22 treated organoids (Fig. 4A). IL-22 is known as a ‘reparative’ cytokine (Lindemans et al. 2015), and the HIPPO pathway is important for regeneration and repair (Gregorieff et al. 2015; Yui et al. 2018), and HIPPO and STAT3 signaling are known to be intertwined (Taniguchi et al. 2015). Thus, it is not surprising to find the HIPPO signature enriched in IL-22 induced genes. However, a link between epithelial IL-13 responses and the HIPPO pathway has not yet been determined. Nonetheless, the large spheroids we observed in IL-13 treated organoids are reminiscent of ‘reparative organoids’ that also grow as large spheroids (Nusse et al. 2018). In addition, activation of the WNT pathway can also lead to spheroid growth (Yin et al. 2014). To test if IL-13 induces spheroids independently of WNT signaling, as suggested by the association with HIPPO signaling, we treated organoids with IL-13, WNT3A, or their combination. Indeed, the combination of IL-13 + WNT3A resulted in a higher proportion of large spheroids than each factor separately (Fig. 4B,C), suggesting that IL-13 induces a WNT-independent growth response. The HIPPO pathway culminates in the activation of its transducers YAP and TAZ. Although, we did not find an increase in *Yap1* expression we did find that the gene encoding TAZ (*Wwtr1*) was markedly increased upon IL-13 treatment (Fig. 4D,E), as well as in vivo after *N. brasiliensis* infection (Fig. 4F). Analogously, upon infection with *N. brasiliensis* we observed an increase in crypt length and enrichment of HIPPO genes by GSEA (Fig. 4G,H). Together, this suggests that IL-13 induces a HIPPO gene expression response *in vitro* and *in vivo*, potentially by controlling TAZ expression.

**Figure 4:**
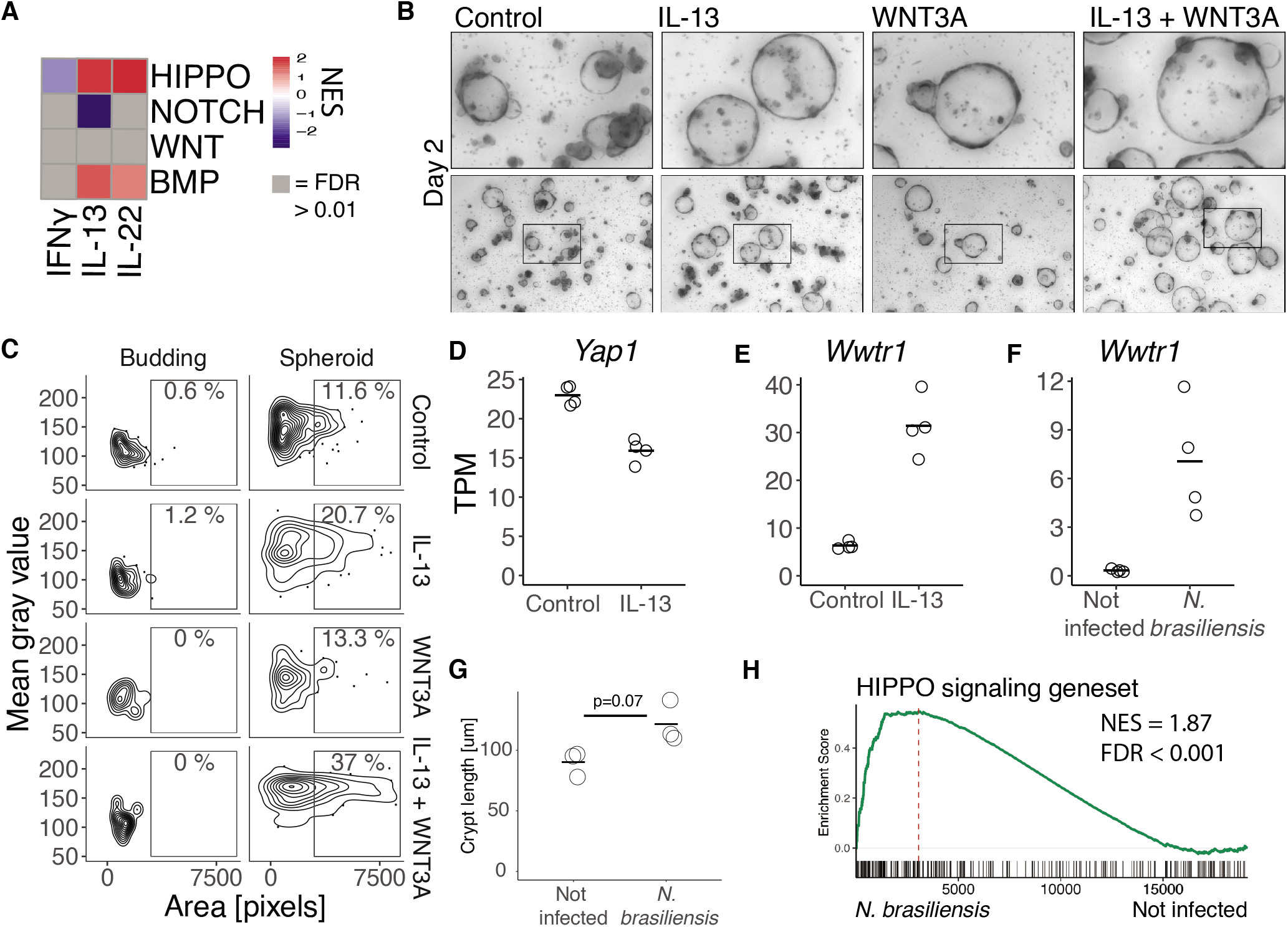
Cytokines have broad effects on signaling pathways that are important in IEC development and homeostasis. **A**, GSEA analysis of gene sets representing developmental pathways important in IEC development of organoids treated with 10 ng/mL cytokine for 24 hours. HIPPO: Genes up in artifical YAP induction and down-regulated in YAP-KO. WNT: Genes up in organoids treated with the GSK3 inhibitor CHIR, NOTCH: Genes up in organoids from ATOH1 KO epithelium compared to wt and BMP: Genes up in organoid treated with BMP4 compared to control. See supplementary file 1 for exact gene sets and references. **B**, Representative images of small intestinal organiods treated with 10 ng/mL IL-13, 25 % media with WNT3A conditioned media or in combination. **C**, Mean gray value and area of organoids from same experiment as B. Representative data of 2 experiments. **D-F**, Expression of indicated genes from RNAseq data, D-E is intestinal organoids treated with cytokines for 24 hours and F is intestinal epithelium infected with *N. brasiliensis.* Adjusted p-values relative to control: *Yap1*: NS, *Wwtr1* (I): < 10^−48^ and *Wwtr1* (J): < 10^−22^. **G**, Quantification of crypt length from cross-sectional sections of intestinal tissue in not infected and mice infected with *N. brasiliensis*. Each circle is the mean of 20 crypts from one biological replicate and statistics determined with two-tailed T-test. **H**, GSEA of a gene set representing HIPPO signaling against RNAseq data from intestinal epithelium from *N. brasiliensis* infection. NES = normalised enrichment Score, TPM = transcripts per million and NS = not significant. All images are projections of Z-stacks.

### BMP signaling is associated with IL-13 and limits tuft cell differentiation *in vitro*

We were somewhat surprised to find enrichment of BMP signaling upon IL-13 treatment (Fig. 4A). BMP members are traditionally expressed by mesenchymal cells, so it is unclear how IL-13 may induce BMP signaling. Nonetheless, established BMP target genes *Id1/2/3* are indeed upregulated specifically after IL-13 treatment (Fig. 5A). Next, we assessed the expression pattern of the TGF-β family members and surprisingly found that IL-13 robustly induced *Bmp2* but not any other members (Fig. 5B, S5A). In support, there was also an increase of *Bmp2* and the TGF-β family member *Inhbc* during a *N. brasiliensis* infection (Fig. 5C,D). Interestingly, Haber et al. lists Bmp2 as a bona fide Tuft cell marker in their gene sets from plate-based scRNAseq of small intestinal epithelium (Haber et al. 2017), and we found an enrichment in Bmp2 expression in Tuft cells (Fig. 5E). The connection between Tuft cells and IL-13 signaling is further highlighted by our finding that tuft cells specifically expressed high levels of *IL13ra1* (Fig. 5F), whereas *IL4ra* or other cytokine receptors did not have such skewed cell-type specific receptor expression (Fig. S6). To test whether *Bmp2* activation is a direct consequence of the IL-13-STAT6 axis, we used a STAT6 ChIP-seq data set (Czimmerer et al. 2018) and found that STAT6 was present near the *Bmp2* transcription start site (TSS) upon IL-4 treatment (Fig. 5G). To test whether BMP signaling affects tuft cell differentiation, we compared organoids grown in normal EGF, NOGGIN, and RSPO (ENR) media with organoids grown in ER media, without the presence of the BMP antagonist NOGGIN. We found that BMP antagonism by NOGGIN is important for tuft cell differentiation as measured by GSEA, established tuft cell marker gene expression, and confocal imaging (Fig. 5H-K).

**Figure 5:**
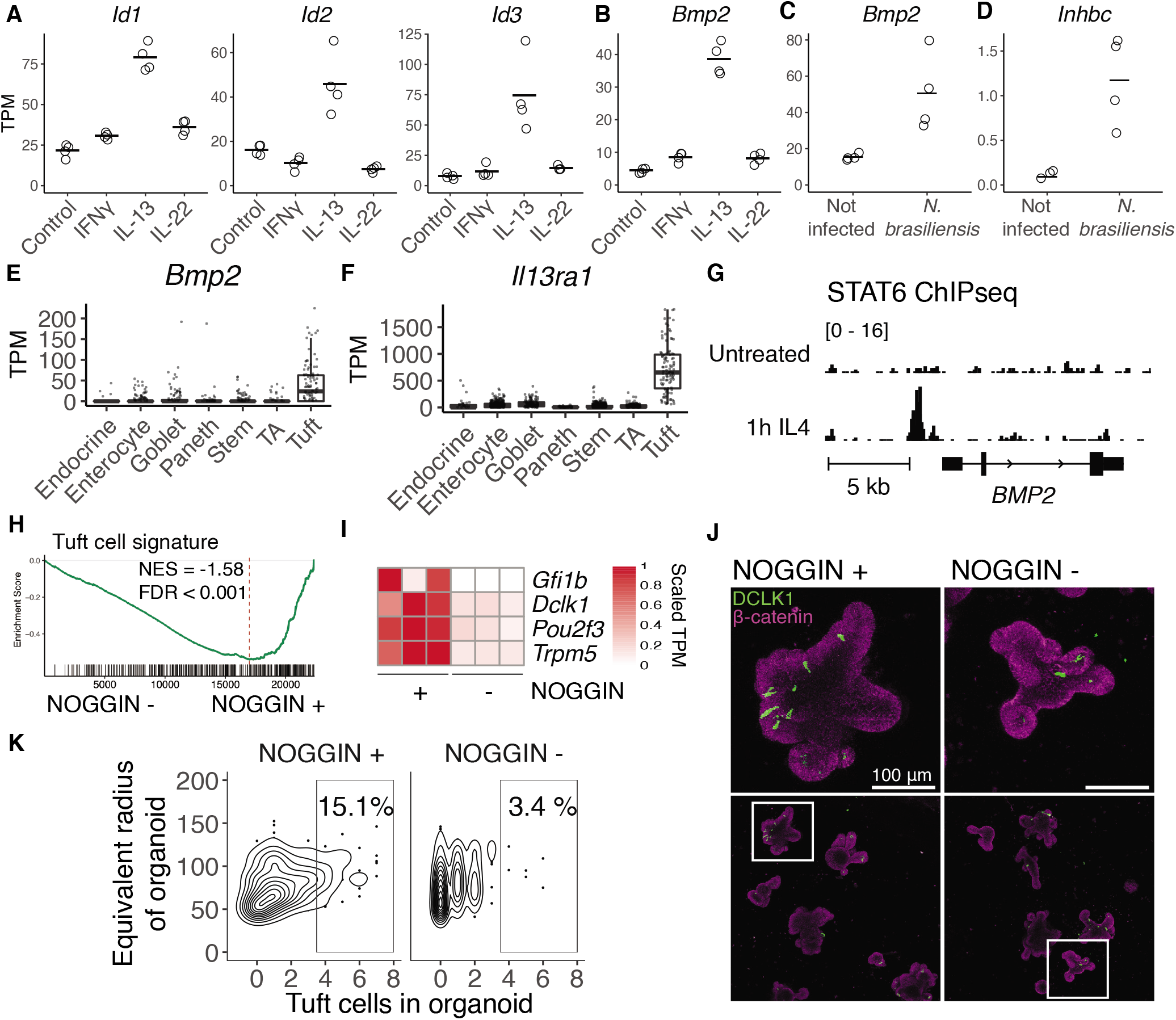
IL-13 induce BMP2 in intestinal epithelium. **A,B**, Gene expression in small intestinal organoids treated with cytokines for 24 hours determined with RNAseq. Adjusted p-values relative to control for IL-13 was < 10^−12^ for all shown genes. **C,D**, Gene expression in intestinal epithelium extracted from mice infected with *N. brasiliensis* determined with RNAseq. Adjusted p-values was < 10^−5^ for *Bmp2* and < 0.001 for *Inhbc.* **E, F**, Plate based scRNAseq expression data from Haber et al in small intestinal epithelium. TA = Transit amplifying. **G**, Stat6 ChIP from human macrophages stimulated with IL4 and control. Shows representative track out of two replicates for each treatment. **H**, GSEA of Tuft cell gene set on RNAseq of small intestinal organoids grown in the presence of NOGGIN or not for 5 days since splitting. **I**, Tuft cell marker genes from same RNAseq data as H. **J**, DCLK1 confocal staining of small intestinal organoids grown in the presence of NOGGIN or not for 72 hours since splitting. **K**, Quantitation of experiment in J. Plot shows a total of 462 organoids. TPM = transcripts per million, NES = normalized enrichment score.

### BMP signaling acts as a feedback loop to limit tuft cell expansion

Tuft cells are crucial mediators of type 2 immunity. In a feed-forward loop, tuft cells amplify ILC2s by expressing IL-25, and ILC2s, in turn, express IL-13 to expand tuft cells (Gerbe et al. 2016; Howitt et al. 2016; Moltke et al. 2016). However, this model lacks any limiting factors that are needed to return back to homeostasis when the infection is cleared. Our findings so far suggest that IL-13 treatment induces an epithelial-intrinsic feedback loop mediated by BMP signaling. To further investigate this, we tested the ALK2 (BMP receptor) inhibitor dorsomorphin homolog 1 (DMH1) in combination with IL-13 (Mohedas et al. 2013). We found that DMH1 did not directly affect IL-13-induced *Bmp2* expression (Fig. 6A), but it did completely block the induction of canonical BMP target genes *Id1/2/3* (Fig. 6B). Furthermore, the enrichment of BMP signaling pathway genes by IL-13 is blocked by DMH1, without modulating the effect of IL-13 on HIPPO or NOTCH pathways genes (Fig. 6C). Although DMH1 did not change the number of tuft cells by itself, it did increase tuft cell numbers upon IL-13 co-treatment (Fig. 6D,E). IL-13+DMH1 also increased expression of tuft cell associated genes such as *Dclk1, Pou2f3, Trpm5*, and *Alox5* (Fig. 6F) as well as the overall tuft cell signature by GSEA (Fig. 6G). Together, this supports a model in which activation of BMP signaling limits IL-13-induced tuft cell differentiation, thus, providing a feedback loop to limit tuft cell expansion and its associated type 2 immunity.

**Figure 6:**
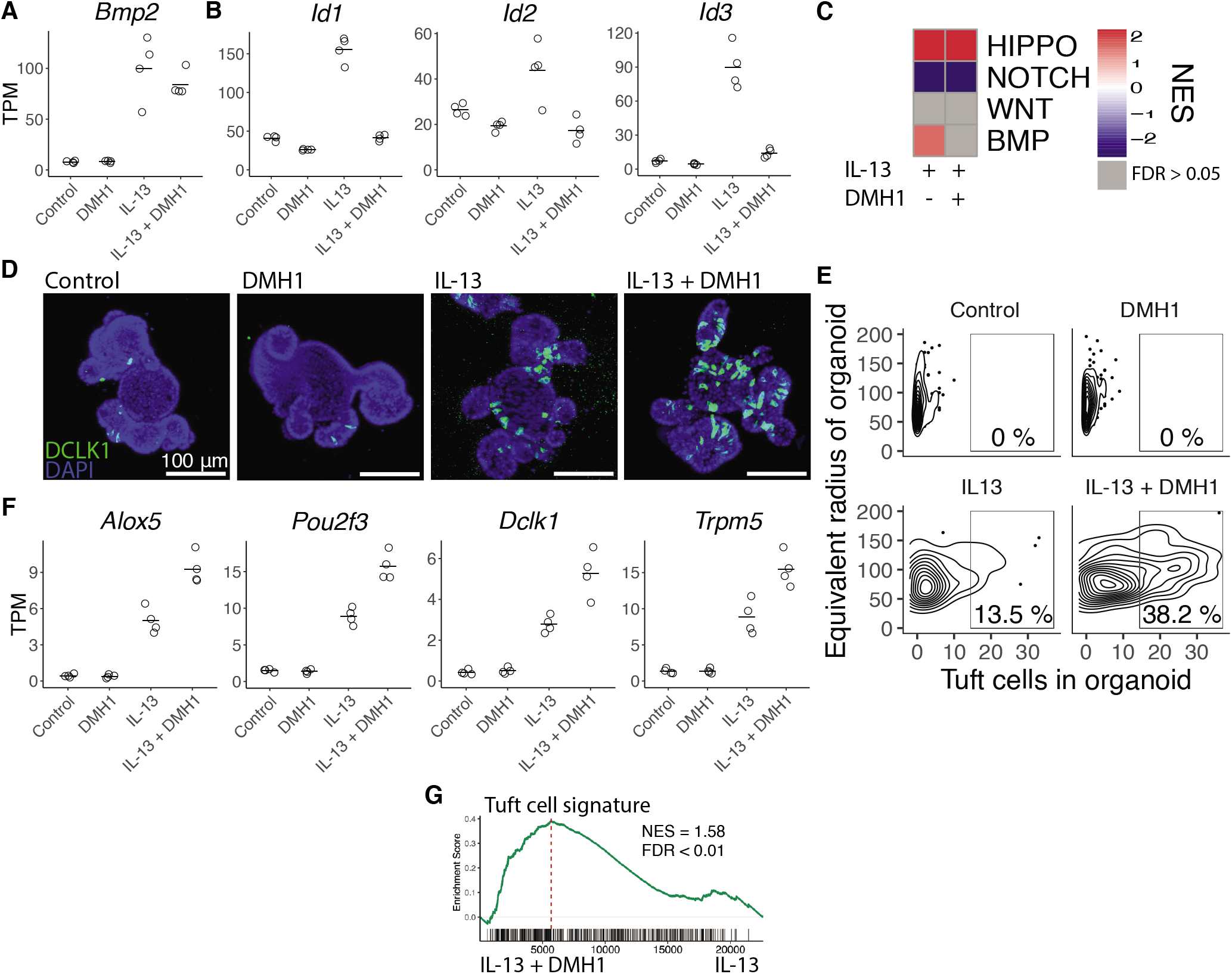
Bmp signaling restricts IL-13-induced tuft cell expansion. **A,B**, Gene expression determined with RNAseq of small intestinal organoids treated with 10 ng/mL IL-13, 5 μM DMH1 or in combination for 72 hours. Each treatment has four biological replicates. Adjusted p-value compared to control: **Bmp2**: DMH1: NS, IL-13: < 10^−68^, IL-13+DMH1: < 10 “A **Id1**: DMH1 < 10^−6^, IL-13:< 10^−49^, IL-13+DMH1: NS, **Id2**: DMH1 < 10 ^−2^, IL-13: < 10 ^−4^, IL-13+DMH1: < 0.05, **Id3**: DMH1 < 0.05, IL-13: < 10 ^−74^, IL-13+DMH1: < 10 ^−4^. **C**, Heatmap of NES values from GSEA of gene sets representing signaling pathways in indicated treatments vs control condition. See supplementary file 1 for exact gene sets. **D**, Confocal staining of DCLK1 in small intestinal organoids treated with indicated treatments for 72 hours. **E**, Quantification of same experiment as D, plot represents a total of 566 organoids. Representative of 4 independent experiments in 4 mice. **F**, Expression of tuft cell markers determined as in A. Adjusted p-values for IL-13+DMH1 compared to control for all genes was < 10^−78^. **G**, GSEA of tuft cell signature in IL13 + DMH1 treated vs IL13 treated.

## Discussion

The intestinal epithelium is capable of rapidly altering its cellular composition to defend against pathogens. Here, we provide a comprehensive comparison of how different immune responses mechanistically drive changes in the intestinal epithelium. Specifically, we find that key cytokines associated with different modes of immunity influence developmental pathways to guide changes in epithelial composition.

A common challenge for studies using intestinal organoids is the large heterogeneity in growth within a single culture well (Merenda, Fenderico, and Maurice 2019). This heterogeneity is readily visible in organoid morphology, and in this study we successfully utilize a neural network to characterize these morphological differences by automatically classifying organoids as “spheroid” or “budding” structures. Despite the biological processes underlying the formation of spheroid and budding organoids not being fully understood (Merenda, Fenderico, and Maurice 2019), these classifications can be used to determine differences between different conditions (Han et al. 2020; Schwank et al. 2013). Furthermore, our quantification of confocal images reveals that the spheroid and budding organoids have different tuft cell numbers, showing that classification is linked to biological implications. We expect that such precise methods for quantification will be important in future work to describe biological mechanisms.

In recent years, tuft cells have taken centre stage in the defense against helminth infections (Schneider, O’Leary, and Locksley 2019). They express receptors that help them detect helminths, which together with type 2 cytokines results in their expansion (Nadjsombati et al. 2018; Schneider, O’Leary, Moltke, et al. 2018). Thus, this culminates in a feed forward loop where tuft cell derived IL-25 activates ILC2s and ILC2 derived IL13 induces tuft cell differentiation (Gerbe et al. 2016; Howitt et al. 2016; Moltke et al. 2016). Importantly, there have been no descriptions of limiting factors in this feed-forward loop, which is obviously required to resolve type 2 immunity. Here, we identify how intestinal-epithelial intrinsic *Bmp* signaling can act as a brake on tuft cell expansion (Fig. 6). In addition, we find that *Il13ra1*, which is involved in responding to type 2 cytokines, is uniquely expressed in tuft cells. We would speculate that this is instrumental for tuft-cell specific signaling responses to type 2 cytokines, for example to induce *Bmp2*.

In addition to our findings regarding cytokine-driven tuft cell expansion, we also provide data suggesting that IL-13 and IL-22 induce different types of goblet cells (Fig. 3). Both cytokines are known to induce goblet cells, and IL-22 is also important for type 2 immunity-driven goblet cell hyperplasia *in vivo* (Turner, Stockinger, and Helmby 2013). Additionally, the goblet-cell effector RELM-beta is critical for resistance against both bacterial and helminth infections (Artis et al. 2004; Bergstrom et al. 2015; Propheter et al. 2017). We hypothesize that the difference is related to the type of immune response. Type 2 immunity requires a ‘weep and sweep’ response, in which goblet cells play an essential role by secreting mucus (weeping) to facilitate the expulsion of helminths (Gause, Rothlin, and Loke 2020), whereas type 3 responses may rely less on the induction of mucus. Instead, we find that IL-22 induces goblet-cell specific ER stress response genes, which corroborates recent work that identified a pathologically relevant role for IL-22 in inducing ER stress response genes (Powell et al. 2020).

Overall, our approach of using intestinal organoid-cytokine co-cultures has revealed that developmental pathways underlie the cellular compositional changes that occur as part of immune responses taking place in the intestine. These changes have been confirmed using relevant *in vivo* models of infection. It is important to balance both active immunity/inflammation with resolution of inflammation, and many immunopathologies reflect the importance of maintaining this balance. The discovery of an innate BMP-driven brake on IL-13 induced immune changes suggest that targeting developmental pathways may therefore be a useful tool to aid resolution of inflammation in clinically relevant diseases.

## Acknowledgment

We thank the animal care (CoMed) and imaging (CMIC) core facilities that assisted in this work (NTNU). Part of the RNA-seq was done by the Genomics Core Facility at NTNU, which receives funding from the Faculty of Medicine and Health Sciences and Central Norway Regional Health Authority. This work was financially supported by the Norwegian Research Council (Centre of Excellence grant 223255/F50, and ‘Young Research Talent’ 274760 to MJO) and the Norwegian Cancer Society (182767 to MJO). This work was supported by the Wellcome Trust through an Investigator Award to RMM (Ref 106122), and the Wellcome Trust core-funded Wellcome Centre for Integrative Parasitology (Ref 104111).

## Code and data availability

- Code for brightfield and confocal analysis: https://github.com/havardtl/coco
- ArrayExpress accession for RNAseq of cytokine treated organoids: E-MTAB-9182
- ArrayExpress accession for RNAseq of *N. brasiliensis*: E-MTAB-9183
- ArrayExpress accession for RNAseq of *C. rodentium*: E-MTAB-9184
- ArrayExpress accession for RNAseq of IL-13 and DMH1 treated organoids: E-MTAB-9185

## Supplementary file consisting of figures S1-S6 and full methods

**Figure S1:**
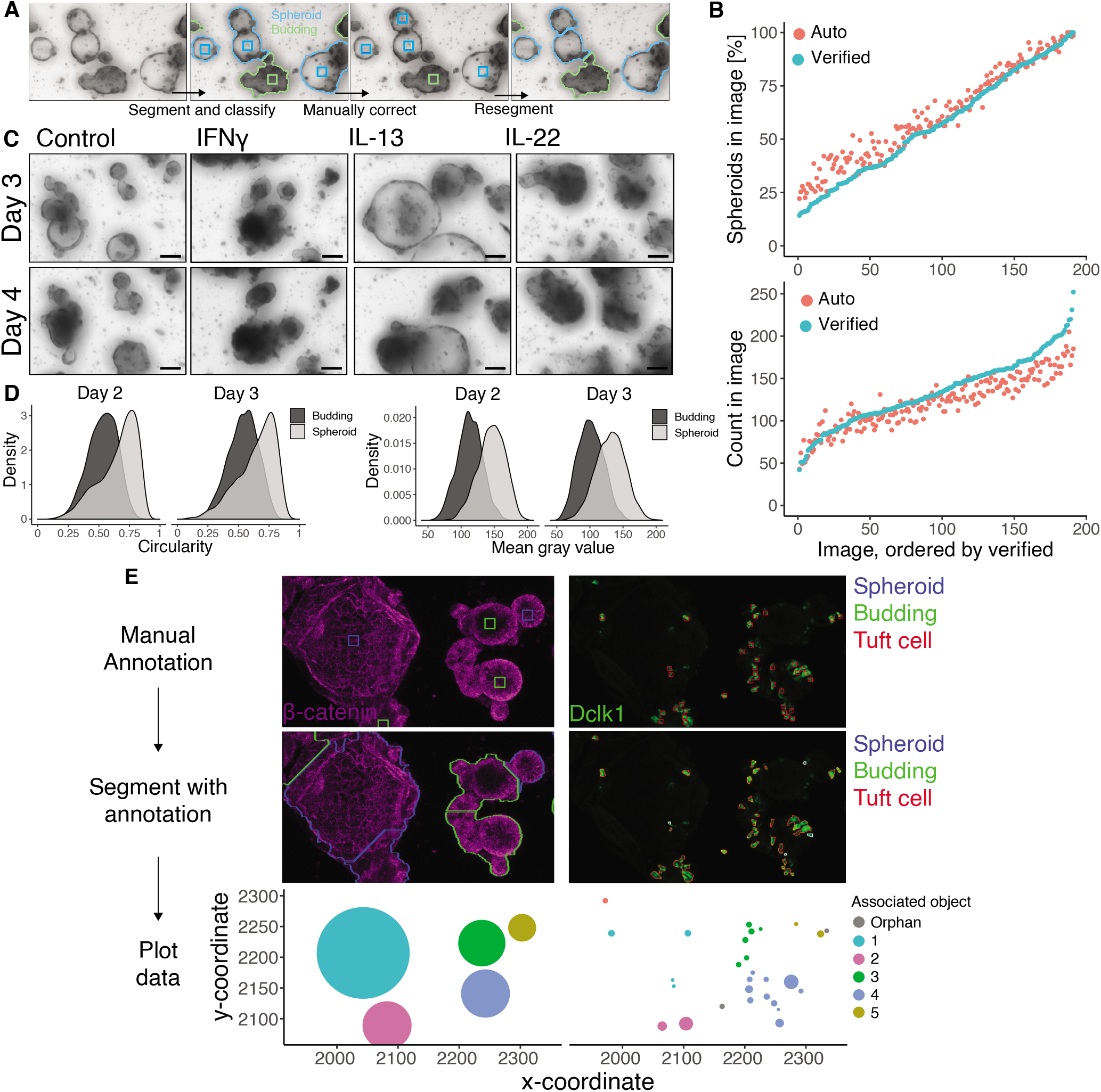
The image analysis pipeline performs well both in segmentation and classification of organoids, and is perfected by a manual verification step. **A**, A segmentation algorithm is used to find the outline and center of an organoid and the segmented organoid is classified by a neural network. An optional manual correction of organoid centers and classification can be performed. The organoids can then be segmented again which gives an updated segmentation following the manual correction. **B**, Comparison of automatic segmentation and automatic classification of organoids compared to manually verified segmentation and manually verified classification. Each circle represents one image. **C**, Minimal projections of small intestinal organoids treated with cytokines. Days since seeding. **D**, Distribution of circularity and mean gray value (higher value is whiter) of untreated small intestinal organoids. **E**, Image output from each step in the confocal image analysis pipeline. The center of each object of interest is marked with a square in a custom graphical user interface, and this information is used to separate even overlapping objects. Furthermore, objects in other channels, like Tuft cells, are analyzed to map which organoid they are inside. Here this association is shown by the organoids and the tuft cells that are within them having the same color in the last panel. This information is summarized and used to plot number of Tuft cells per organoid.

**Figure S2:**
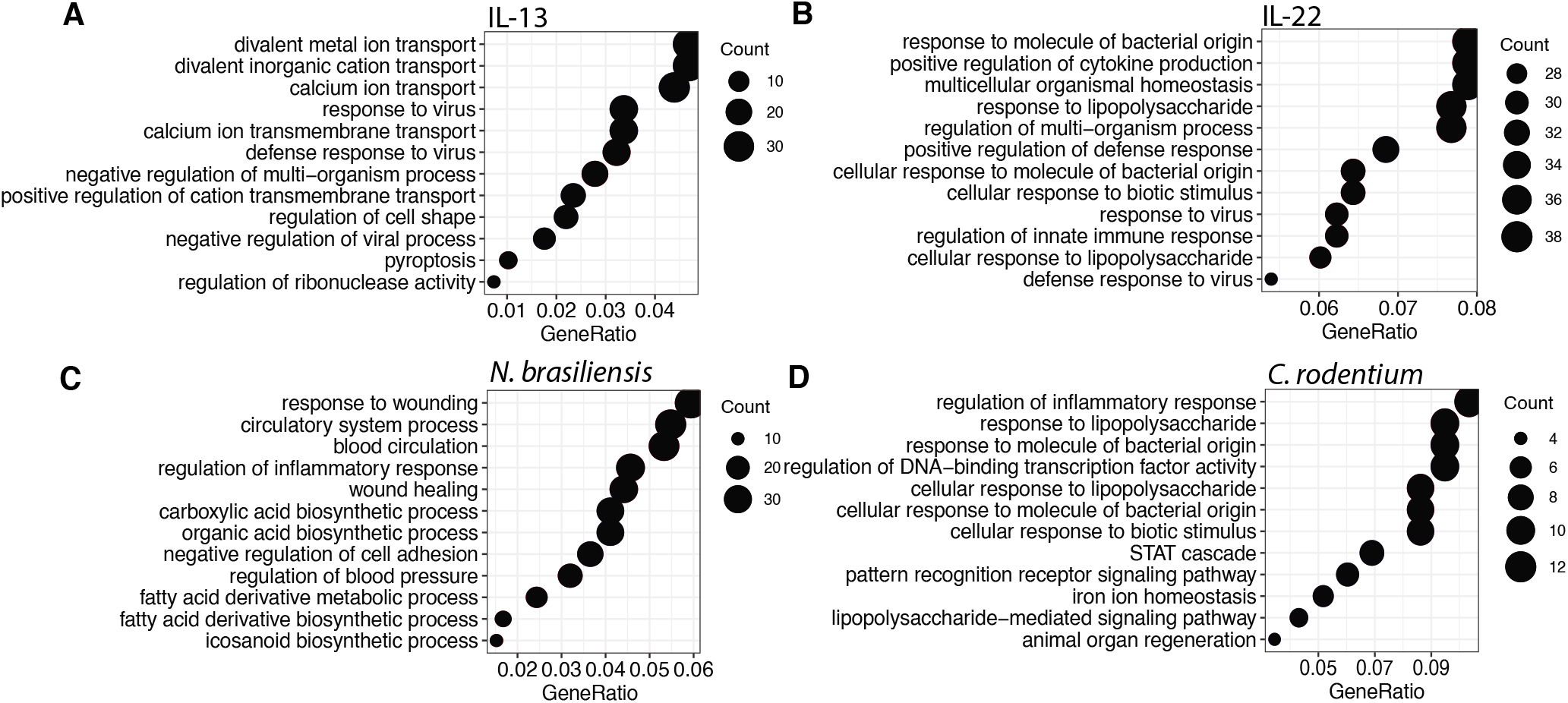
Details of RNA sequencing data and flow cytometry. **A-D**, Showing top hits from GO-term analysis of “biological process” in indicated RNAseq data.

**Figure S3:**
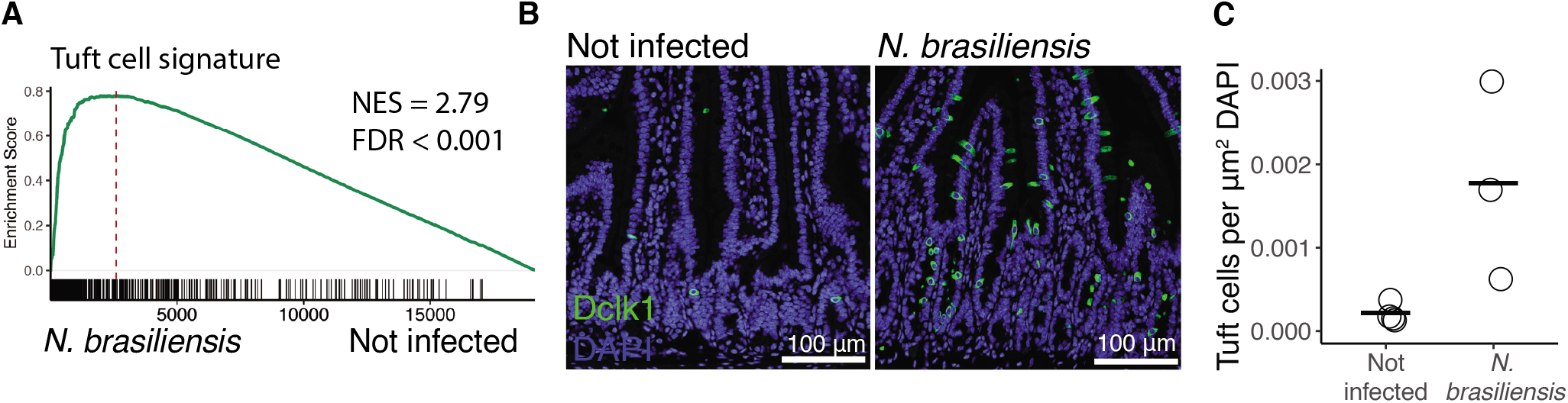
*N. brasiliensis* infection induce tuft cells. **A**, GSEA analysis of a tuft cell signature on mRNA data from intestinal epithelium from *N. brasiliensis* infected mice. **B**, Confocal microscopy of cross section of intestine from mice infected with *N. brasiliensis* and not infected. **C**, Automatic Tuft cell count from same experiment as B. Each circle represents quantitation from one full cross section from one mouse.

**Figure S4:**
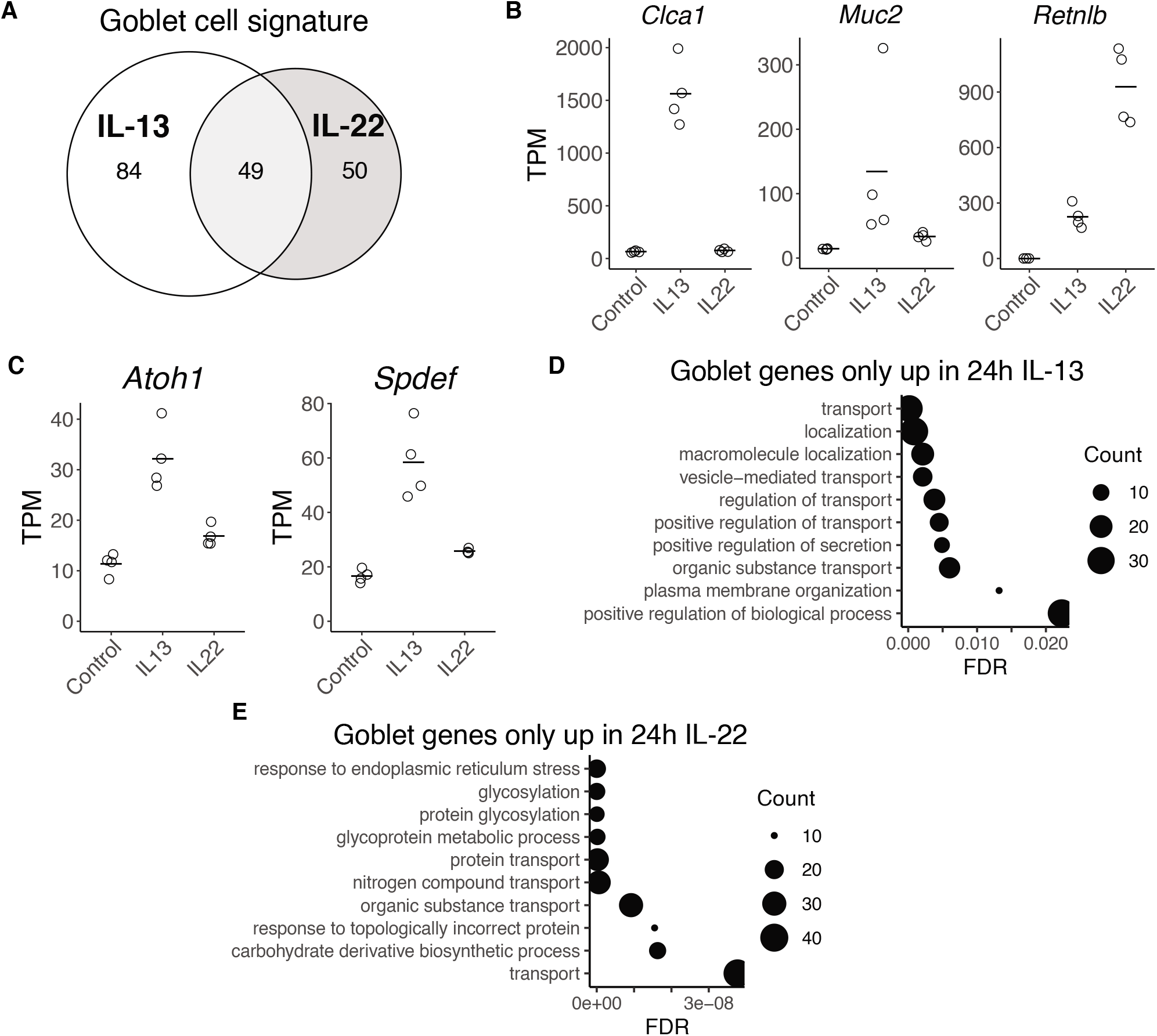
IL-13 and IL-22 gives different goblet cells also after 72 hours of cytokine stimulation. **A**, Distribution of how the plate based scRNAseq goblet cell gene set from Haber et al is changed upon 72 hours of IL-13 and IL-22 treatment. Up is defined as lg2fc>0.5 and p-adj<0.05. **B,C**, Gene expression from intestinal organoids treated for 72 hours with indicated cytokine. Adjusted p-values relative to control: *Clca1*: IL-13: < 10^−168^, IL-22: < 0.1, *Muc2*: IL-13: < 10^−6^ IL-22: < 10^−26^, *Retnlb*: IL-13: < 10^−62^, IL-22: < 10^−84^. *Atoh1*: IL-13: < 10^−17^, IL-22: < 10^−3^, *Spdef*: IL-13 < 10^−21^, IL-22: < 10^−8^. **D,E**, GO terms associated with gene sets seen in venndiagram in figure 3B from the online STRING tool. FDR = False discovery rate.

**Figure S5:**
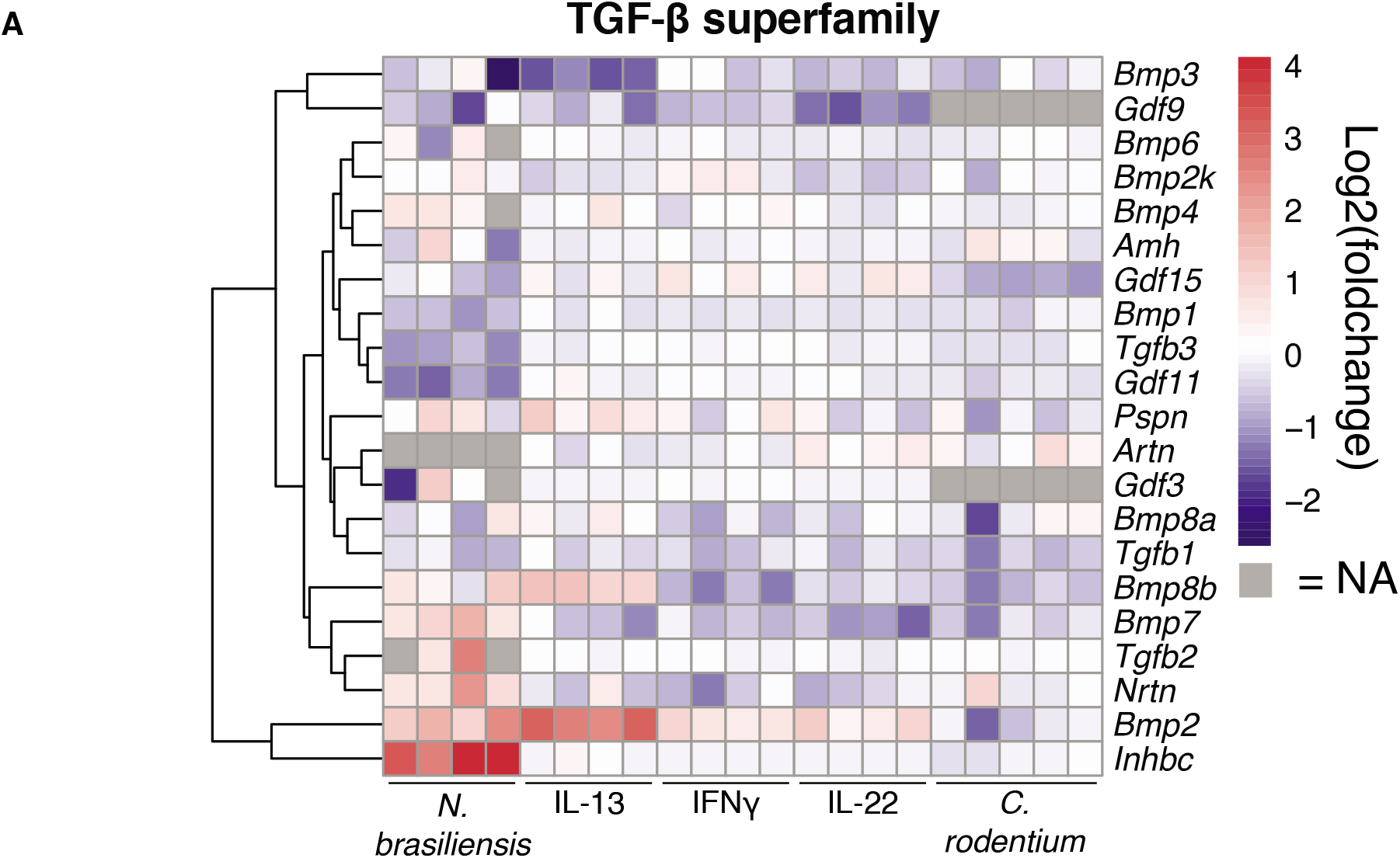
BMP2 is the main TGF-β superfamily member induced by IL-13. **A**, Heatmap of log2 fold change for TGF-β superfamily members from mRNAseq data from intestinal epithelium from mice infected with *N. brasiliensis* or *C. rodentium* and organoids treated for 24 hours with indicated cytokine.

**Figure S6:**
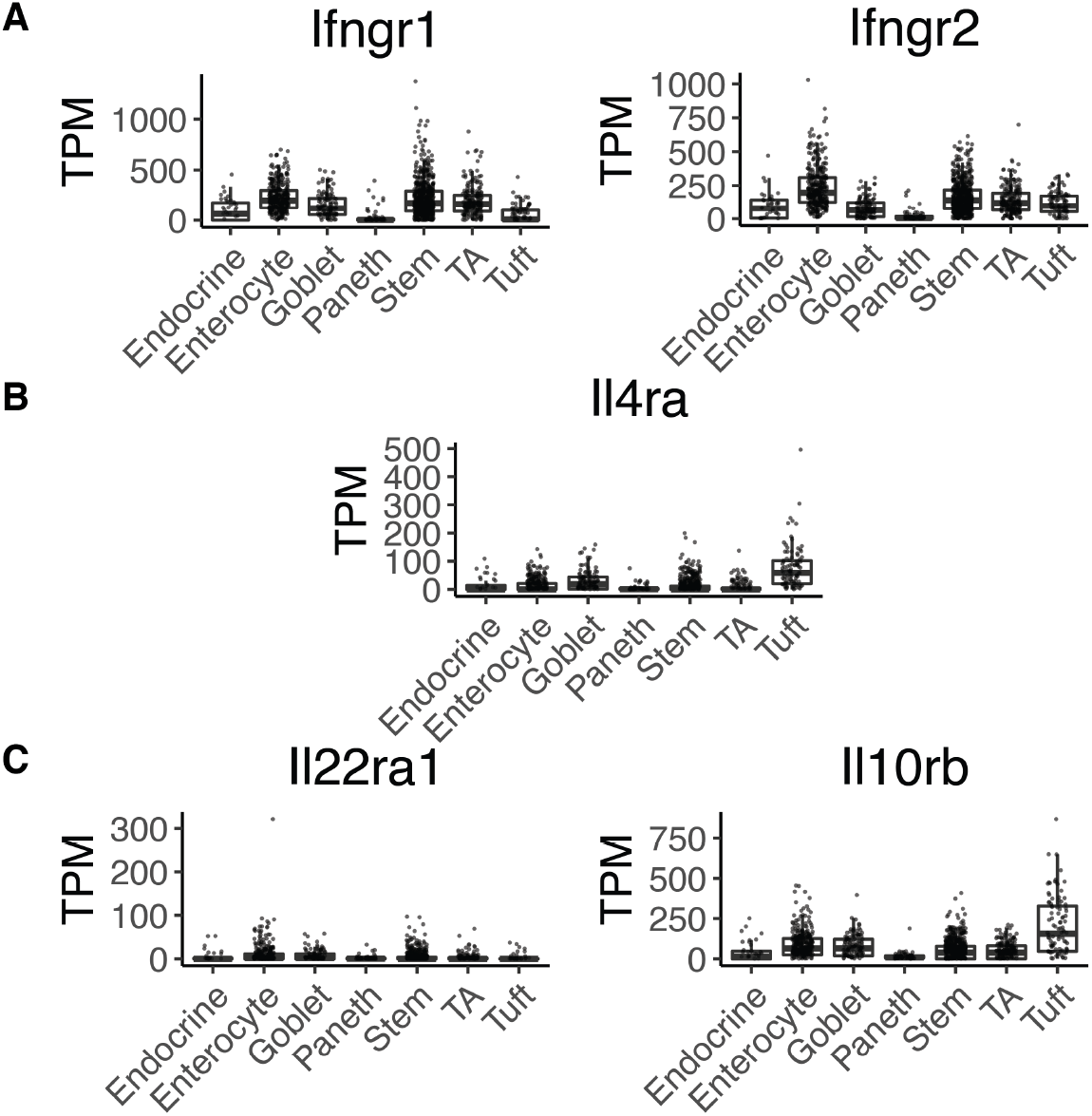
Cell specific expression of cytokine receptors. **A-C**, Expression of cytokine receptors, for IFNγ (A), IL-13 (B) and IL-22 (C), from plate based scRNAseq from Haber et al. TA = Transit amplifying.

## Methods

### Animal experiments

All mouse work done in Norway was performed in accordance with CoMed NTNU institutional guidelines and Norwegian legislation on animal experiments. Mice were handled under pathogen-free conditions. The C57BL/6 mice strain was used for all experiments.

The mice studies for *Citrobacter rodentium* infection were approved by Norwegian Food Safety Authority (FOTS ID: 11842). Briefly, C. rodentium was grown at 37 °C in Luria-Bertani (LB) medium supplemented with chloramphenicol (30 μg/ml) overnight. 9-11 weeks old mice were orally gavaged with a culture of 10^8^ – 10^9^ CFU per mouse delivered in a volume of 100 μL of sterile PBS. Mice remained infected with *C. rodentium* for six days. To determine the infection level, fecal samples were collected on day 5, homogenized in sterile PBS with a FastPrep homogenizer. After serial dilution of fecal samples, homogenates were plated on chloramphenicol resistant agar plates and the plates were counted after 24 h of growth at 37 °C. After 6 days of infection, mice were euthanized by CO_2_ inhalation, parts of distal colon tissue was harvested for isolation of crypts as described below. RNA was isolated and samples were further used for RNAseq analysis.

Mice studies with *Nippostrongylus brasiliensis* infection used mice aged 8-12 weeks maintained in individually ventilated cages according to UK Home Office guidelines. *N. brasiliensis* was maintained as previously described (Camberis, Le Gros, and Urban 2003). Infections were carried out using 200 L3 *N. brasiliensis* larvae administered subcutaneously. After 7 days of infection, mice were euthanised via cervical dislocation, and duodenal tissue harvested to use in imaging and for isolation of crypts as described below.

### Small intestine and colon crypt isolation

Crypts from small intestine and colon where isolated according to a published protocol (Sato and Clevers 2013). In summary, the small intestine or distal colon of mouse was washed in ice-cold PBS and opened longitudinally. A microscopy cover slip was used to gently scrape of excess mucus and villus before the intestine was cut into 2-4 mm pieces. The fragments were resuspended in 30 mL ice-cold PBS and pipetted up and down with a 10 mL pipette. Supernatant was discarded and this step was repeated 5-10 times until the supernatant was clear. The fragments were incubated in 2 mM EDTA in PBS for small intestine, and 10 mM EDTA in PBS for colon, at 4 °C for 30 min with gentle rocking. The supernatant was removed, 20 mL of ice-cold PBS was added and the fragments were pipetted up and down. This washing step was repeated until the crypt fraction appeared as seen in a microscope. The 1-3 consecutive crypt fractions were passed through a 70 μm cell strainer and collected into a FCS coated 50 mL tube. The crypts were spun down at 300 x*g* for 5 min and washed once in 10 mL ice cold PBS at 200 x*g* for 5 min.

### Small intestinal organoid culture

Approximately 200-500 crypts were resuspended in 50 μL Matrigel (Corning, 734-1101) and kept at 4 °C. 50 μL of the matrigel solution was added to the center of a pre-warmed 24 well plate or 8-well microscopy slide (Ibidi, 80821) and quickly transferred to an incubator at 37 degrees with 5 % CO_2_. After 5 min the pellet had solidified and 500 μL of basal culture medium was added and the plate was put back into the incubator. Basal culture medium consisted of advanced Dulbecco’s modified Eagle medium - F12 supplemented with penicillin/streptomycin, 10 mM HEPES, 2 mM Glutamax, 1x N2 [ThermoFisher Scientific 100X, 17502048], 1x B-27 [ThermoFisher Scientific 50X, 17504044], and 1x N-acetyl-L-cysteine [Sigma, A7250]) and overlaid with ENR factors containing 50 ng/ml of murine EGF [Thermo Fisher Scientific, PMG8041], 20 % condition medium (CM) from a cell line producing R-Spondin (kind gift from Calvin Kuo, Stanford University School of Medicine, Stanford, CA, USA) and 10 % Noggin-CM (a kind gift from Hans Clevers, Hubrecht Institute, Utrecht, The Netherlands). The culture medium was replaced every 2-3 days. Organoids were passaged by disrupting them with strong mechanical pipetting, letting the solution cool on ice before centrifuging at 200 x*g*, 5 min at 4 °C and resuspending in matrigel.

### Bright field imaging and quantification of intestinal organoids

Z-stacks covering the entire matrigel droplet were captured using a EVOS2 microscope with CO2, temperature and a humidity-controlled incubation chamber (Thermo-Fisher Scientific). 2D morphological properties of organoid objects as well as their classification were gathered using a custom analysis program written in python based on opencv2 (Bradski 2000). The brightness of images were autoscaled to max brightness, and a canny edge detection algorithm was run on each individual z-plane using the opencv2.canny function. Groups of pixels below a certain size was removed and a minimal projection of the edges was generated. This image was used to define the contour of objects. The implementation of opencv2s wathershed algorithm was used to split somewhat overlapping objects from each other and the center of the object was defined as the pixel furthest from the edge of the object. Each defined object contour was used to extract a 120×120 pixel picture of the object on a white background from a minimal projection of the original z-stack. These images were used to classify the organoid as either “Junk”, “Budding” or “Spheroid” with a convolutional neural network implemented using Tensorflow and Keras (Abadi et al. 2015; *Keras* 2015). The network was trained on about 25 000 manually classified 120×120 images prepared as just described. For initial analysis these data were plotted, but for plots used in this publication the pictures were reviewed manually in a GUI written in python. This GUI displayed the center of each object and its classification and enabled addition of new object centers or editing of automatically found object centers. The manually verified annotations were used to re-run the segmentation as described above, but this time the classification of the objects were not changed and a watershed algorithm were used to segment object contours based on the manually curated object centers. Finally, objects with a size less than 300 were filtered out of the data and plots were made with the R-package ggplot2 (Wickham 2016).

### Immunofluorescence staining of organoids and imaging

The matrigel used for organoids grown for immunofluorescent imaging were cultured in matrigel mixed with 25 % BCM in eight-well slides (Ibidi, 80821). Media was removed and organoids were fixed in 300 μL 4% PFA with 2 % sucrose for 30 minutes. Fixation solution was removed and 300 μL PBS was added, after 5 minutes the PBS was removed. This washing step was done a total of two times. The PBS was removed and 0.2% Triton X-100 in PBS was added for 30 minutes at room temperature to permeabilize the cell membranes. The wells were then washed in PBS, 3 times 5 minutes each time. Free aldehydes were blocked in 100 mM Glycine for 1 hour at room temperature and thereafter the wells were incubated in 300 μL blocking buffer (2 % normal goat serum, 1 % BSA and 0.2 % Triton X-100 in PBS) for 1 hour at room temperature. Primary antibodies, β-catenin (1:200, mouse mAb, Santa Cruz Biotechnology, sc-7963), DCLK1 (1:250, rabbit pAb, abcam, ab31704), MUC2 (1:200, rabbit pAb, Santa Cruz Biotechnology, sc-7963) and RELM-beta (1:200, rabbit pAb, PeproTech 500-P215) were diluted in 150 μL blocking buffer and incubated overnight at 4 °C. The next day samples were washed in PBS with agitation, 3 times 10 minutes each time. The appropiate Alexa fluor Secondary antibody (1:500), and DAPI (1:10 000) and UEA1 (Ulex Europaeus Agglutinin I Rhodamine-labeled Dil 1:500, Vector laboratories RL1062) was added in blocking buffer (1 % normal goat serum, 0.5 % BSA and 0.2 % Triton X-100 in PBS) and incubated at 4 °C with agitation over night. The next day the samples were washed in PBS, 3 times 5 minutes each time, and 250 μL Fluoromount G medium (ThermoFisher Scientific, 00-4958-02) was added to the well. Z-stacks of tile-scans covering the entire well were acquired on a Zeiss Airyscan confocal microscope using a 10X objective lens. The confocal images of organoids stained with a RELM-beta antibody used a concentration of 5 ng/mL IL-22 but where otherwise treated as the other images. The images stained with RELM-beta antibody where also used in the quantification in another paper (Parmar et al. 2020).

### Immunofluorescence staining of tissue sections and imaging

The harvested intestinal tissue was fixed with 10% formalin for 48 h at room temperature. Tissue was subsequently dehydrated through a series of graded ethanol and then embedded in paraffin wax blocks. 5 μm thick sections were cut using a microtome, floated in a 40 °C water bath and transferred to a glass slide. After drying, the tissue sections were kept in an oven at 60 °C for 30 minutes and deparaffinized in two changes of Neo-clear (Xylene substitute) for 5 min each, followed by graded ethanol (100 % ethanol 2 times with 3 min each, and next transfer once through 95 %, 80 %, and 70 % ethanol and water 3 min each). Heat mediated antigen retrieval was performed using citrate buffer. Tissue sections on glass slides were marked with hydrophobic pen (PAP pen, ab2601) to keep staining reagents on the tissue section. Next, blocking buffer (1% BSA, 2% normal goat serum, 0.2% Triton X-100 in PBS) was added onto the sections of the slides and incubated in a humidified chamber at room temperature for 1 hour. Appropriately diluted primary antibody (antibody dilution buffer, e.g. 0.5 % bovine serum albumin, 1 % normal goat serum, 0.05 % Tween 20 in PBS) was added to the sections on the slides and incubate in a humidified chamber at 4 °C overnight. The slides were washed in 0.2 % Triton X-100 10 min each. Next, secondary antibody conjugated with fluorochromes was added to the slides for 1 hour at room temperature in a humidified chamber. Hoechst was used as counterstain and UEA1 was added to stain goblet cells. After rinsing three times with PBST (0.2 % Triton X-100 in PBS) for 10 min each, slides were mounted in Fluoromount G. Complete tile scans were acquired with a Zeiss Airyscan confocal microscope, using a 20x objective lens.

### Quantification of immunofluorescence images

2D morphological properties of intestinal organoid objects in confocal images were gathered using a custom analysis program written in python based on opencv2 (Bradski 2000). Maximal projections of each channel were created as well as a rgb composite of all channels, hereafter referred to as the combined channel. The center position of all intestinal organoids as well as their class as either “budding” or “spheroid” were manually annotated in the combined channel using a custom graphical user interface. A binary threshold individual to each channel was then applied to create channel masks and additionally, these masks were added together to a combined mask. Contours of objects were defined based on the outline in combined mask and they were split using a watershed algorithm where the manual annotations were used as input. Only objects in the combined channel with a manual annotation were kept for further analysis. 2D morphological properties such as area, perimeter and circularity of each object in the combined channel was acquired as well as the mean intensity of each channel. Furthermore, tuft cells were manually annotated in the DCLK1 channel using the custom graphical user interface. The number of tuft cells inside each object in the combined channel could therefore be counted. These data were processed in R and plotted with the ggplot2 package (Wickham 2016).

The same analysis were applied to confocal images of tissue staining with some changes. A binary mask defining the complete tissue area to quantify, in this case villus and crypt, was manually made for each tissue section. Furthermore, manual annotation of tuft cells was done for a selection of images using a custom written graphical user interface. The number of manual tuft cells inside the manually made tissue section mask was counted. In addition, both the mean intensity of each channel and the number of pixels above a channel-individual threshold was measured for each channel. This enabled a calculation of number of manually counted Tuft cells per μm^2^ DAPI positive pixel. Furthermore, a binary threshold of the DCLK1 channel allowed definition of automatically defined tuft cell objects. Whether a DCLK1 object was a tuft cell or noise could be confirmed by checking for presence of a manual tuft cell annotation. In this way, gates for selecting tuft cells where defined and used for images that were not manually annotated. Counts for manual annotation and automatic annotation were found to correlate closely and automatic annotations are used in the plots in the paper. These data were processed in R and plotted with the ggplot2 package (Wickham 2016).

### Flow cytometry of intestinal organoids

Cytokines were applied to organoids 24 hours after seeding and harvested after 96 hours. A single cell solution was yielded by mechanical disruption of Matrigel, incubation with TrypLE and occasional pipetting. Cells were washed and transferred to 96-well V-bottom plates, stained for 20 min at 4 °C in PBS/2% FCS in presence of Fc receptor blocking reagent (Biolegend) with anti-murine CD326-BV605, I-A/I-E FITC (all Biolegend, 1:200 dilution) and H-2Kb APC (eBioscience, 1:200 dilution), washed, stained with DAPI (1:1000 dilution) for live-dead exclusion. All centrifugation steps were carried out at 300-370x g. Samples and compensation controls were filtered through a 100 μM mesh and immediately analyzed on a LSRII instrument (BD). Samples were analyzed in FlowJo v10.

### RNA isolation and sequencing

RNA was extracted with Quick-RNA Microprep Kit according to the manufacturers instructions (Zymo, R1050). For small intestinal organoids treated with cytokines for 24 hours, organoids where grown for 24 hours without cytokines before 10 ng/mL of cytokine was applied. Library preparation and sequencing for these samples was performed by NTNU Genomic Core facility. Library preparation was done with Lexogen SENSE mRNA kit and the library was sequenced on two Illumina NS500 HO flowcells, 75 bP single stranded. For small intestinal organoids treated with 10 ng/mL cytokines and 5 μM DMH1 for 72 hours, cytokines and DMH1 was applied immediately after seeding. All other library preparation and sequencing was performed by Novogene (UK) Co. using the NEB Next^®^ Ultra™ RNA Library Prep Kit. Samples were sequenced at 150 bp paired end using a Novaseq 6000 (Illumina).

### RNAseq analysis

Reads were aligned to the *Mus Musculus* genome build mm10 using the STAR aligner (Dobin et al. 2013). The count of reads that aligned to each exon region of a gene in GENCODE annotation M18 of the mouse genome (Frankish et al. 2019) was counted using featureCounts (Liao, Smyth, and Shi 2014). Genes with a total count less than 10 across all samples were filtered out. A differential expression analysis was done with the R-package DESeq2 (Love, Huber, and Anders 2014), and volcano plots were plotted with the R-package EnhancedVolcano (Blighe 2019). PCA analysis was performed with the scikit-learn package with the function sklearn.decomposition.PCA (Pedregosa et al. 2011). GSEA analysis was run with the log2(fold change) calculated by DESeq2 as weights, 10000 permutations and otherwise default settings using the R-package clusterProfiler (Yu et al. 2012). The R-packages pheatmap and eulerr where used to make heatmaps and venndiagrams, respectively (Kolde 2019; Larsson 2019). STRING analysis was done by submitting lists of genes to the online STRING database v11.0, disconnected nodes where hidden and edges show confidence based on the string interaction sources: “Experiments”, “Databases”, “Co-expression” and “Co-occurence” (Szklarczyk et al. 2019). RNAseq of organoids grown in the absence and presence of Noggin can be found from the ArrayExpress accession E-MTAB-9181 and analysed as described in (Alonso et al. 2020).

### Chipseq analysis

Raw reads where downloaded from the Sequence Read Archive from accession GSE106706 using default settings with the tool fasterq-dump from sra-toolkit (*SRA-Tools* n.d.). Reads were deduplicated using clumpify.sh from BBmap with settings dedupe=t and subs=2 (Bushnell 2014) and aligned to the *Mus musculus* genome build mm10 using the STAR aligner (Dobin et al. 2013). BAM files were sorted, indexed and counted with samtools (Li et al. 2009). The bamCoverage function in deepTools was used to create bigwig files that were scaled to the sample with the lowest count (Ramírez et al. 2016). Integrative genomics viewer was used to create plots of chipseq tracks (Robinson et al. 2011).

## Supplementary Tables

### Supplementary file 1 - GSEA gene sets

A file with all gene sets used for GSEA. See file *Supplementary_file_1_gsea.xlsx*.

### Supplementary file 2 - Differential expression

A file with differential expression results from relevant comparisons from mRNAseq. The last columns in each sheet (after “padj”) is TPM values for the indicated sample. See file *Supplementary_file_2_diff_expr.xlsx*

